# Magnetic Resonance Spectroscopy as a Non-invasive Method to Quantify Muscle Carnosine in Humans: a Comprehensive Validity Assessment

**DOI:** 10.1101/568923

**Authors:** Vinicius da Eira Silva, Vitor de Salles Painelli, Samuel Katsuyuki Shinjo, Wagner Ribeiro Pereira, Eduardo Maffud Cilli, Craig Sale, Bruno Gualano, Maria Concepción Otaduy, Guilherme Giannini Artioli

## Abstract

Carnosine is a dipeptide abundantly found in human skeletal muscle, cardiac muscle and neuronal cells having numerous properties that confers performance enhancing effects, as well as a wide-range of potential therapeutic applications. A reliable and valid method for tissue carnosine quantification is crucial for advancing the knowledge on biological processes involved with carnosine metabolism. In this regard, proton magnetic resonance spectroscopy (1H-MRS) has been used as a non-invasive alternative to quantify carnosine in human skeletal muscle. However, carnosine quantification by 1H-MRS has some potential limitations that warrant a thorough experimental examination of its validity. The present investigation examined the reliability, accuracy and sensitivity for the determination of muscle carnosine in humans using *in vitro* and *in vivo* experiments and comparing it to reference method for carnosine quantification (high-performance liquid chromatography – HPLC). We used *in vitro* 1H-MRS to verify signal linearity and possible noise sources. Carnosine was determined in the m. gastrocnemius by 1H-MRS and HPLC to compare signal quality and convergent validity. 1H-MRS showed adequate discriminant validity, but limited reliability and poor agreement with a reference method. Low signal amplitude, low signal-to-noise ratio, and voxel repositioning are major sources of error.

## Introduction

Carnosine is a multifunctional dipeptide abundantly found in human skeletal muscle, cardiac muscle, and in some neuronal cells [1]. Carnosine has numerous properties that confers performance enhancing effects [2], as well as a wide-range of potential therapeutic applications [3, 4]. Such properties include hydrogen cation (H^+^) buffering [5], scavenging of reactive species [6], and protection against glycation end products [7]. Several studies have demonstrated the beneficial effects of increased muscle carnosine content (for a comprehensive review, see 1), which can be easily achieved via dietary supplementation of β-alanine, the rate-limiting precursor of carnosine synthesis [8].

A reliable and valid method for tissue carnosine quantification is crucial for advancing the knowledge on biological processes involved with carnosine metabolism, including whether its properties translate into relevant roles for normal physiological function and disease prevention. In human skeletal muscle, carnosine has been quantified in biopsy samples followed by chromatography [8, 9] or mass-spectrometry [6]. Even though obtaining muscle biopsies is a relatively simple and safe procedure [10], the invasive nature of the muscle biopsy technique limits its application, which warrants the development of valid and reliable non-invasive alternatives.

In this regard, proton magnetic resonance spectroscopy (1H-MRS) has been used as a non-invasive alternative to quantify carnosine in human skeletal muscle [11-13]. In 1H-MRS, carnosine is quantified from two detectable signals emitted by the carbon four (C4-H) and the carbon two (C2-H) of the imidazole ring, which resonate at seven and eight ppm of the magnetic resonance spectrum [14]. Although 1H-MRS has been used to quantify carnosine in numerous investigations, there has not been any comprehensive investigation of the validity of 1H-MRS for muscle carnosine assessment against a reference method, such as the chromatographic determination in muscle biopsy samples.

Carnosine quantification by 1H-MRS has some potential limitations that warrant a thorough experimental examination. Firstly, *in vivo* carnosine signals are broad, of small amplitude, often close to the noise level, and tend to suffer dipolar coupling, in particular the signal emitted by C4-H [14]. This makes C4-H quantification unfeasible in most cases [14]. Secondly, the *in vivo* spectrum is crowded with metabolite peaks, thereby making carnosine identification particularly challenging, even when prior knowledge-based approaches are used [15, 16]. Thus, carnosine quantification appears to be more difficult than other abundant muscle metabolites, such as creatine, taurine and lactate [17]. Thirdly, the signals emitted by the imidazole ring could, in theory, also be detected in other imidazole-containing molecules, such as free imidazole, free histidine, carnosine analogues and histidine residues in proteins. In fact, previous investigations have reported problems in differentiating signals from carnosine and its analogue homocarnosine in human brain [18]. This could represent a confounding factor for carnosine quantification by 1H-MRS. Fourthly, carnosine concentrations are not homogenous in muscle tissue, since fibre type distribution may affect local carnosine concentrations [9, 19]. Also, fat tissue surrounding the measurement area can suppress the 1H-MRS signal, adding another source of variation to the carnosine signal [14-16]. Lastly, other variables, such as the signal-to-noise ratio, different angle and/or site of quantification, different machine operators and different data treatment can have major influences on metabolite quantification [14-16, 20, 21].

To address the potential limitations of 1H-MRS to quantify carnosine in human skeletal muscle, the present investigation examined the reliability, accuracy and sensitivity of 1H-MRS for the determination of muscle carnosine in humans using *in vitro* and *in vivo* experiments. Carnosine determination by high-performance liquid chromatography (HPLC) in extracts from human muscle biopsy samples was used as the reference method.

## Results

### 1H-MRS loses signal quality when measuring muscle carnosine in-vivo

To examine whether signal quality is affected when carnosine is measured *in vivo* (*i.e.*, matrix effect), the signal obtained from a pure carnosine solution (12.5 mmol·L^-1^ carnosine phantom solidified in 2% agarose) was compared with the signal obtained *in vivo* from human calf muscle displaying similar carnosine concentration (10.4 mmol·L^-1^). Pure carnosine phantom showed excellent signal quality, with sharp peaks and high signal-to-noise ratio (figure 1, upper panel), whereas the signal obtained *in vivo* showed broadened peaks of lower intensity and lower signal-to-noise ratio (figure 1, lower panel).

**Figure 1.**
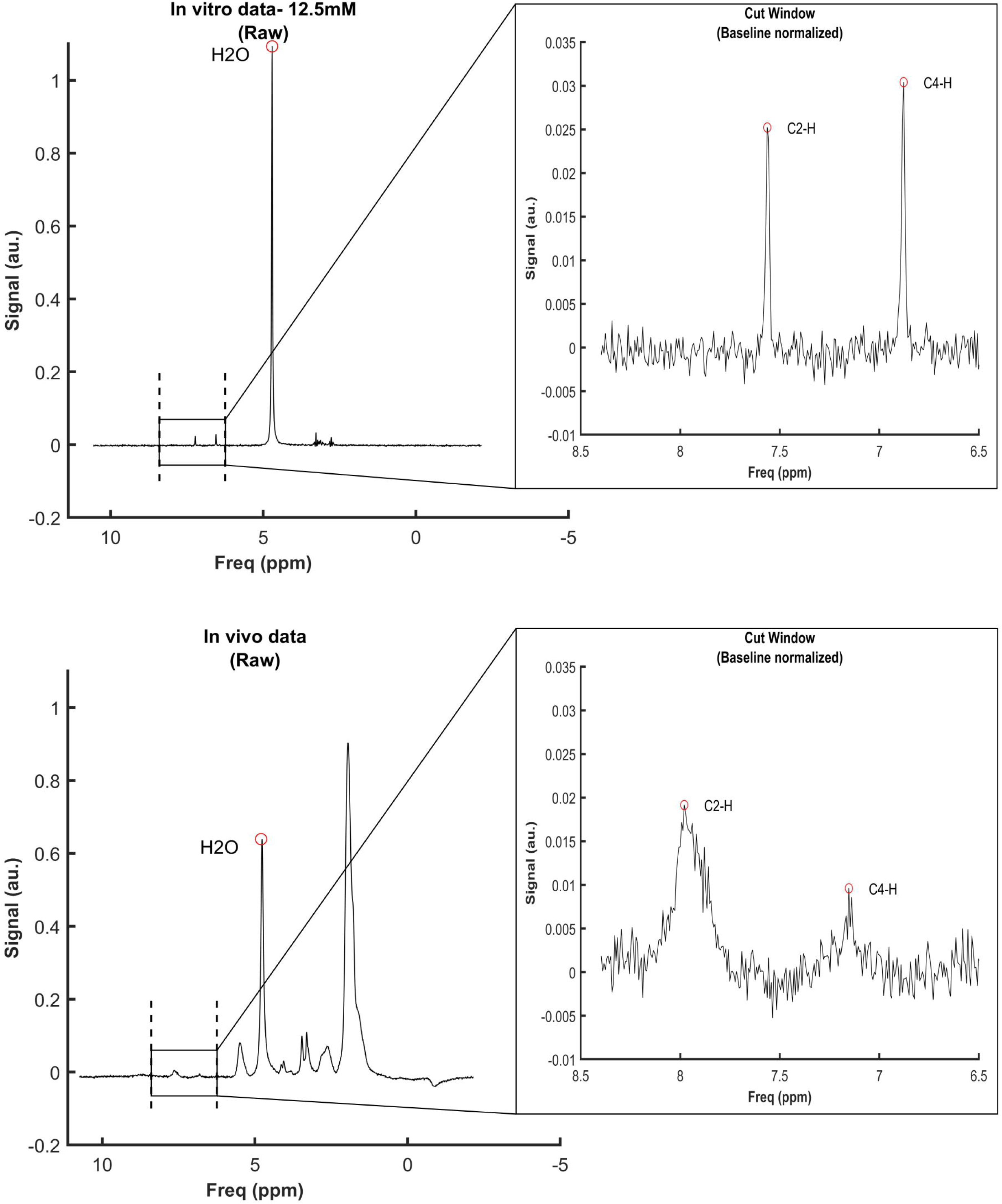
Signal obtained from a phantom containing a 12 mmol·L^-1^ solidified pure carnosine solution (top panel) and signal obtained *in vivo* from the *m. gastrocnemius* of a similar carnosine concentration (10.4 mmol·L^-1^) (bottom panel). On the left, the signals are plotted as raw data; water peak is circled and the frequency area containing the carnosine peaks is indicated by the vertical dashed lines. On the right, the cut window of the frequency area containing the carnosine peaks are shown, indicating the signal after normalization.

### 1H-MRS detects signal emitted by small, but not large, imidazole-containing molecules

To examine the potential interference of other sources of imidazole ring to the carnosine signal obtained *in vivo*, we performed a series of *in vitro* 1H-MRS acquisitions in phantoms containing pure imidazole (12.5 mmol·L^-1^), pure histidine (equivalent to imidazole 12.5 mmol·L^-1^), pure carnosine (equivalent to imidazole 12.5 mmol·L^-1^) and pure protein (bovine serum albumin, BSA, equivalent to imidazole 12.5 mmol·L^-1^).

Signals were quantifiable in the phantoms containing imidazole, histidine and carnosine, but not in the phantom containing protein (figure 2), indicating that small molecules containing imidazole are potential interferences in carnosine quantification, whereas histidine residues in large proteins are not. The carnosine signal was sharp and of low noise, whilst imidazole and histidine signals were less sharp and had larger noise in relation to their amplitude.

**Figure 2.**
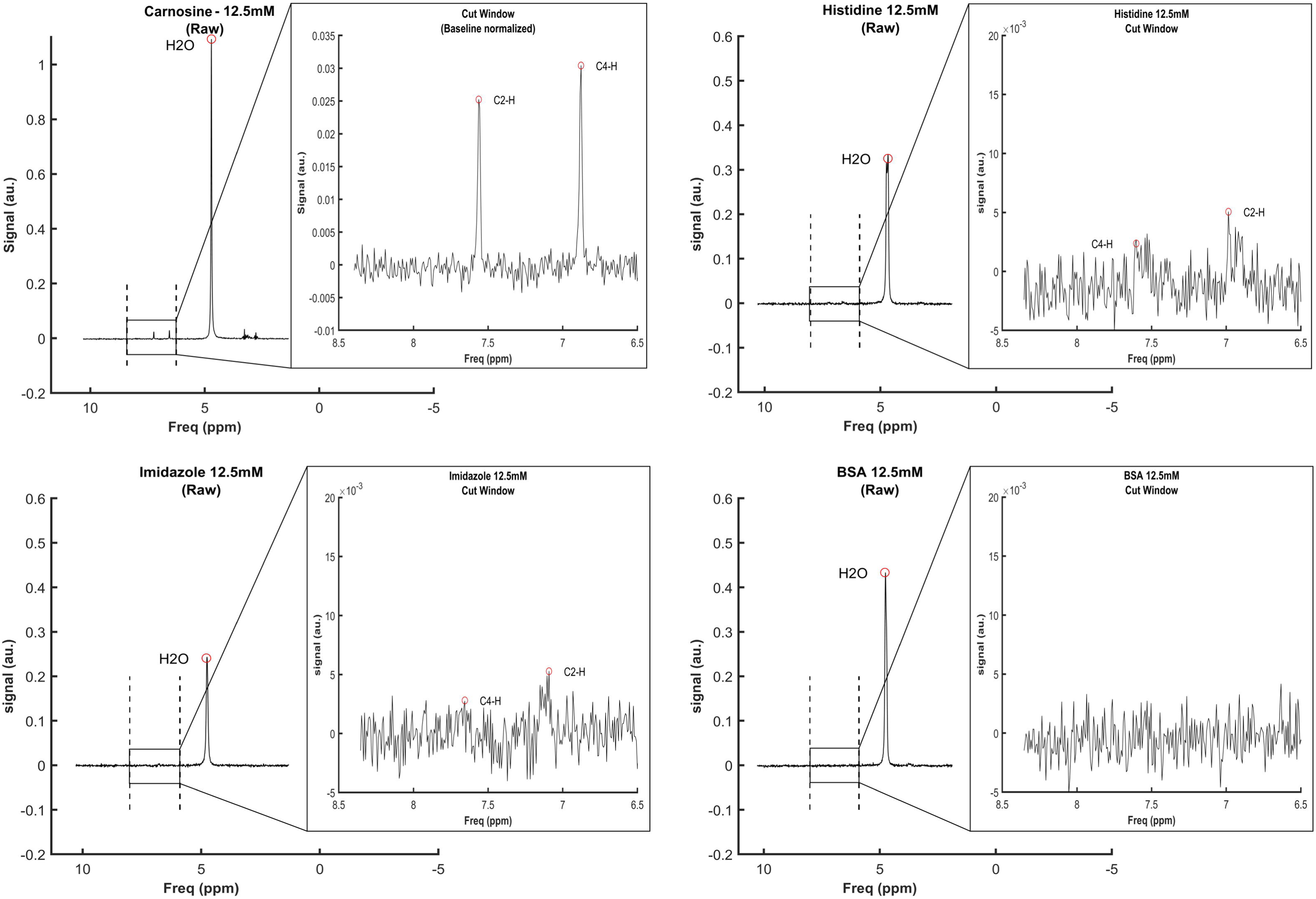
Signal linearity obtained from phantoms containing different concentrations of imidazole, histidine and carnosine. Concentrations are equimolar in imidazole.

### 1H-MRS displays excellent signal linearity within the physiological concentration range

To assess signal linearity of small imidazole-containing molecules, a series of acquisitions were performed in phantoms containing different concentrations of pure imidazole (12.5, 25.0 and 50.0 mmol·L^-1^), pure carnosine (equivalent to 3.0, 4.5, 6.0, 12.5, 25.0 and 50.0 mmol·L^-1^ of imidazole) and pure histidine (equivalent to 12.5, 25.0 and 50.0 mmol·L^-1^ of imidazole). Excellent signal linearity was shown for all substances within the concentration range assayed (figure 3).

**Figure 3.**
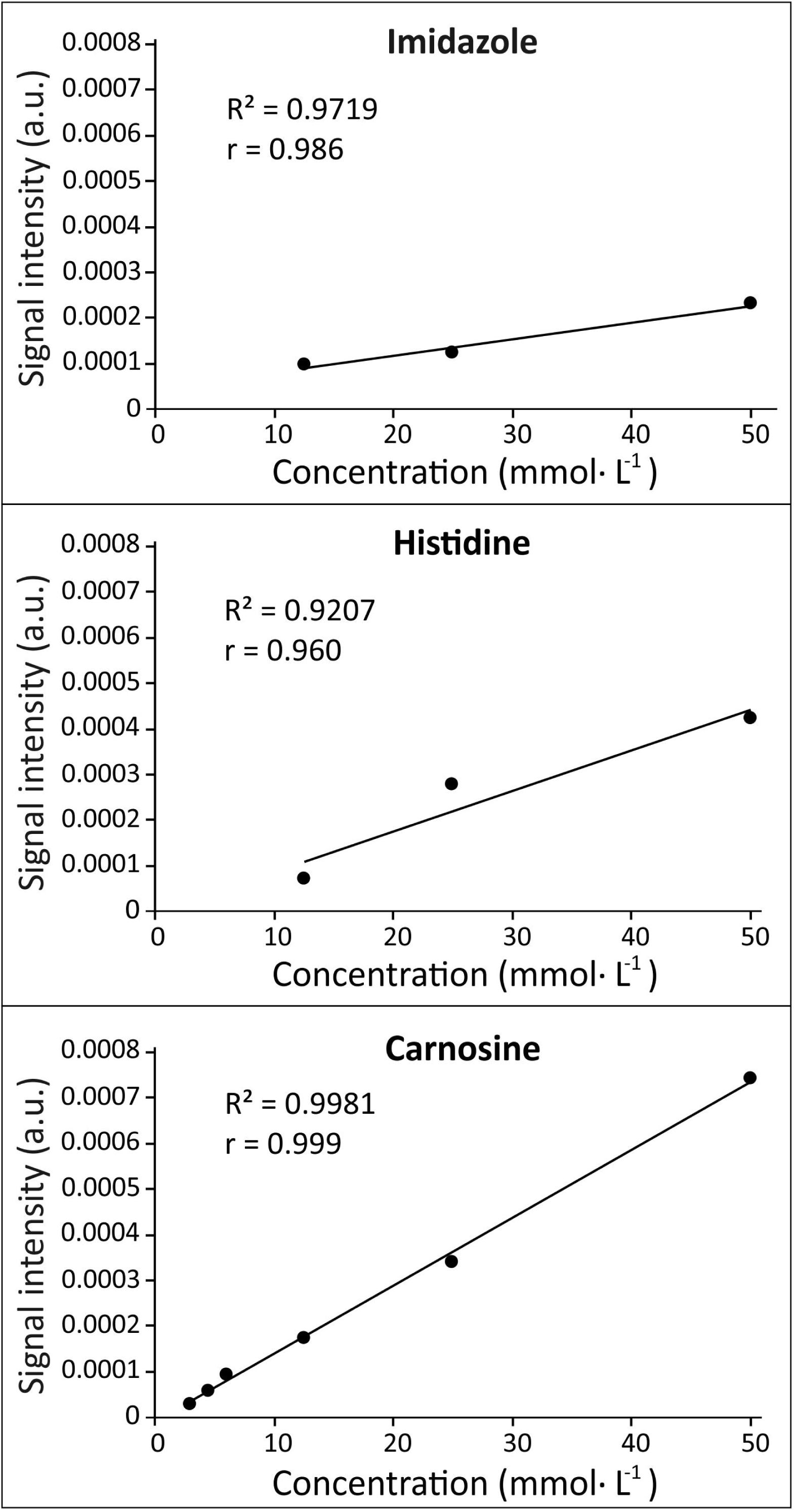
Spectra obtained from phantoms containing 12.5 mmol·L^-1^ of carnosine, histidine and imidazole (panels A, B and C, respectively) and bovine serum albumin (BSA) (panel D). Signals were quantifiable in all spectra, except BSA.

### HPLC as a reference method

In order to confirm that HPLC could be used as a reference method, and to account for all major sources of error associated with this method, we determined intra-assay reliability (*i.e.*, same extract, from the same muscle sample, analysed on two separated runs; n=15 *m. vastus lateralis* samples), inter-extract reliability (*i.e.*, two different extracts, from the same muscle sample, analysed on two separated runs; n=11 *m. vastus lateralis* samples) and inter-biopsy reliability (two different extracts from two different biopsies of the same muscle; n=7 *m. gastrocnemius* samples).

Intra-assay reliability showed very similar values between measurements (mean difference: 0.6±4.0%). No statistically significant differences between measurements were shown (t=0.144; p=0.887), indicating that HPLC is free of systematic errors. A high ICC (0.996, 95%CI=0.987-0.999) and low CV (2.7%) were shown between measurements (Figure 4, panel A). Inter-extract reliability also showed very similar values between measurements (mean difference: 1.0±4.5%), with no statistically significant differences between them (t=0.519; p=0.615). A high ICC (0.988, 95%CI=0.956-0.997) and low CV (3.2%) were shown between measurements (Figure 4, panel B). Inter-biopsy reliability analysis also showed very similar values between measurements (mean difference=1.1±6.0%) with no statistically significant differences between them (t=-0.588; p=0.578). A high ICC (ICC=0.957, 95%CI=0.750-0.993) and low CV (3.9%) were shown between measurements (Figure 4, panel C).

**Figure 4.**
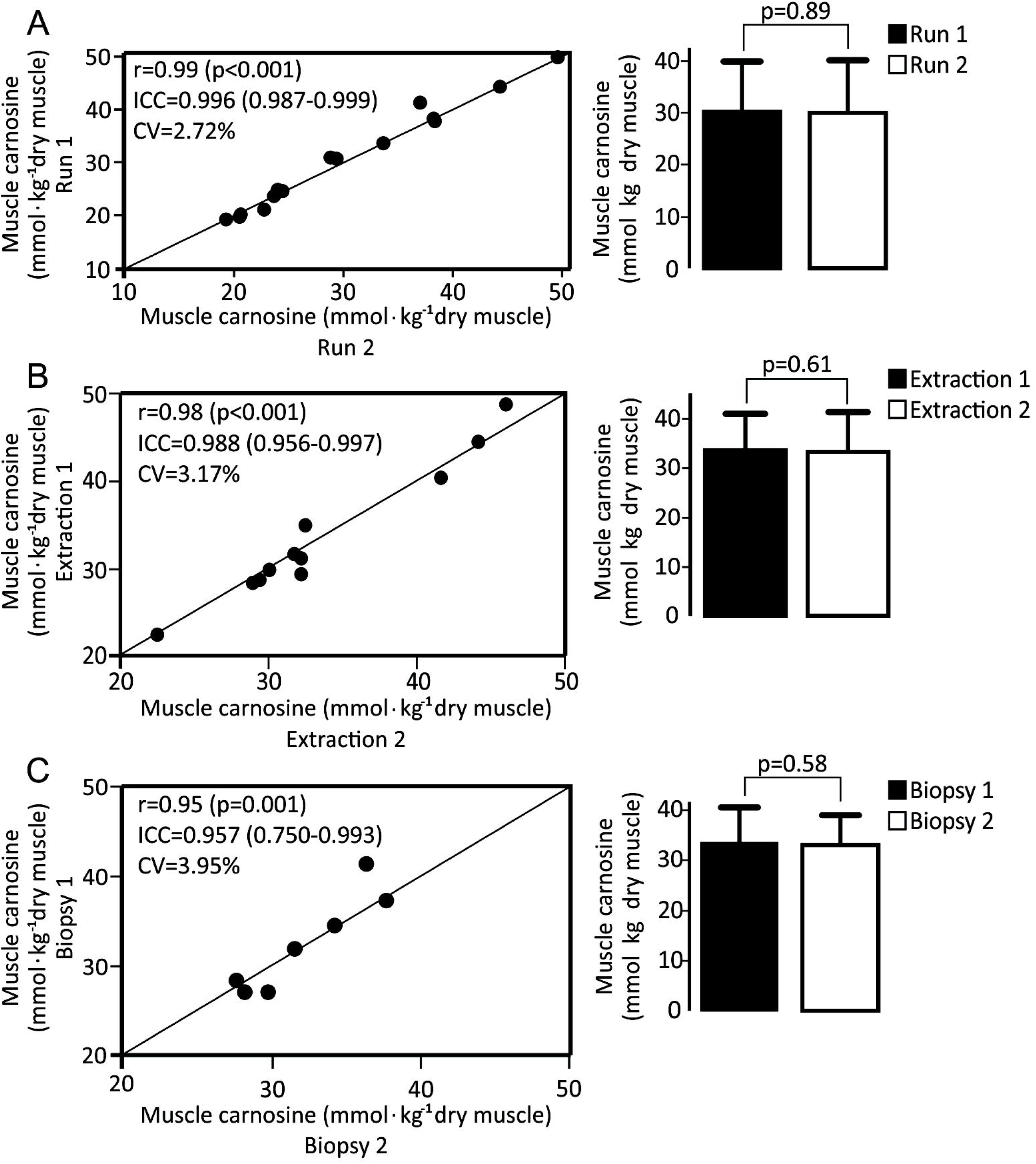
Repeatability analysis of the chromatographic determination of carnosine in muscle samples considering the intra-assay variability (same extraction from the same samples measured twice – panel A), “inter-extract” variability (different extractions from the same samples, measured in duplicate – panel B) and “inter-biopsy” variability (different extractions from different samples, measured in duplicate – panel C). The left charts depict test-retest agreement for each individual sample (in comparison with the line of identity) along with indexes of reliability. The right charts depict the mean ±SD for test and retest conditions.

### Validity assessment of 1H-MRS to quantify carnosine in-vivo in human skeletal muscle

To thoroughly assess the validity of 1H-MRS to quantify carnosine in human skeletal muscle, a series of *in-vivo* studies were conducted in young healthy men aiming to examine test-retest reliability, discriminant validity (*i.e.*, ability to differentiate conditions of knowingly different carnosine concentrations) and convergent validity (*i.e.*, agreement with quantification via HPLC in extracts obtained from muscle biopsy samples).

#### Poor reliability of 1H-MRS to quantify muscle carnosine

1H-MRS test-retest reliability was assessed in two different conditions: with (n=10) and without (n=5) the participant being removed from the scanner and the voxel being repositioned and re-shimmed before the second test. When the participants were not removed from the scanner, good reliability was obtained between measurements, (ICC=0.924, 95%CI=0.451-0.992; CV=6.6%; Figure 5, upper panel). However, reliability indexes were poorer when individuals were removed and subsequently repositioned on the equipment, and the voxel was repositioned and re-shimmed (ICC=0.775, 95%CI=0.325-0.939; CV=16.9%; Figure 5, lower panel). No statistically significant differences were shown between tests in both conditions, indicating that 1H-MRS is free of systematic errors.

**Figure 5.**
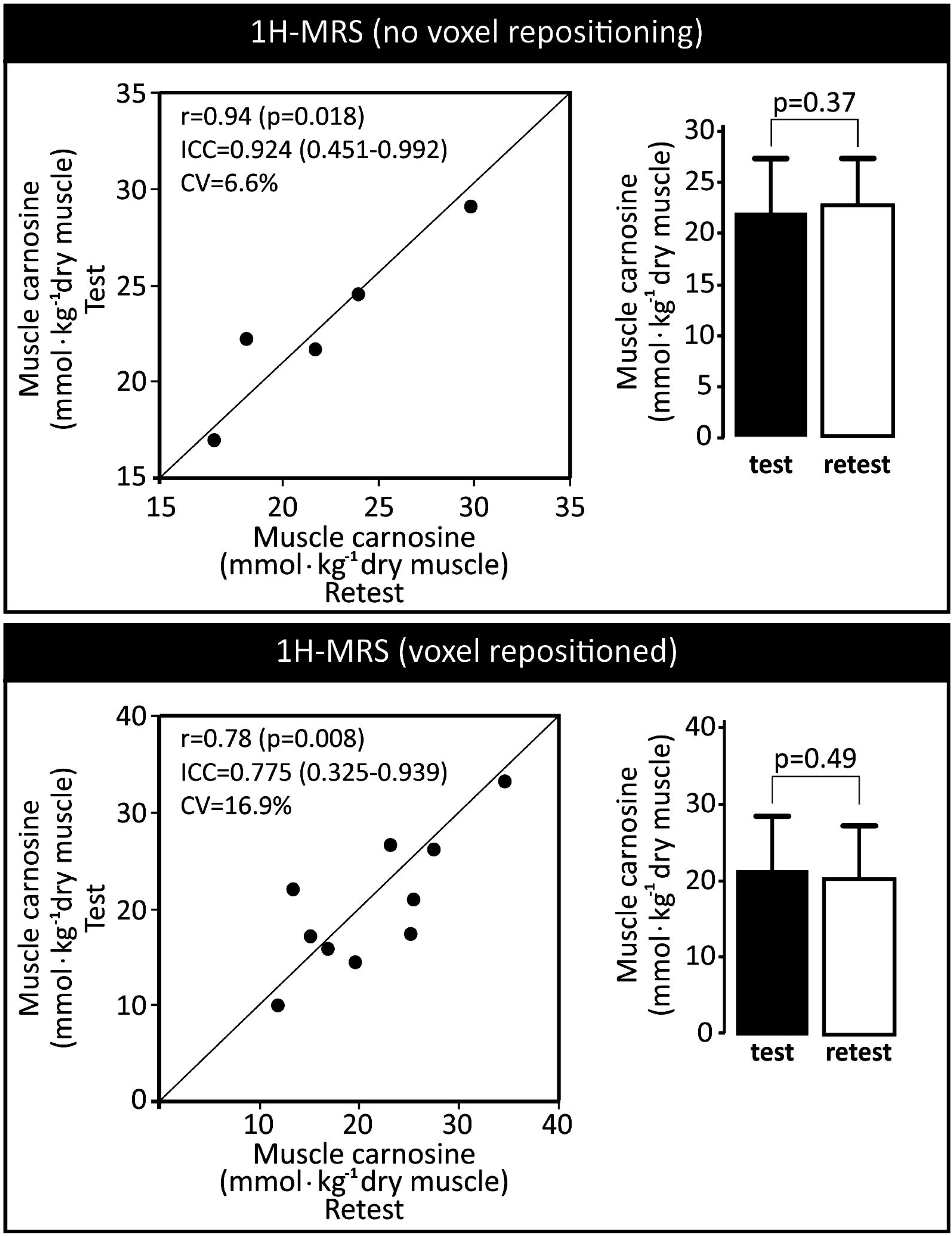
Repeatability analysis of the *in vivo* 1H-MRS determination of carnosine. In the upper panel, test and retest were undertaken without removing the participant from the scanner and without repositioning and replacing the voxel. In the lower panel, the participants were removed from the scanner after test and the retest was undertaken with the voxel being repositioned and re-shimmed. The left charts depict test-retest agreement for each individual test (in comparison with the line of identity) along with indices of reliability. The right charts depict mean ± SD for test and retest.

#### 1H-MRS has adequate discriminant, but poor convergent validity to quantify muscle carnosine

Muscle carnosine was determined using both 1H-MRS and HPLC in the *m. gastrocnemius* of 14 participants before and after 4 weeks of β-alanine supplementation. Both methods were able to detect a significant increase in muscle carnosine after β-alanine supplementation (figure 6, panel A). When comparing the absolute and relative delta increase in muscle carnosine measured by both methods, no statistically significant differences were shown for the mean values between methods (figure 6, panel B), therefore indicating that 1H-MRS has adequate discriminant validity. However, when individual data were analysed and delta changes (either absolute or percent changes) obtained with 1H-MRS are plotted against HPLC, a large disagreement was shown, and most of the measurements were further away far from the line of identity (figure 6, panels C and D). This is confirmed by the Bland-Altman plot including the entire data set, which shows a large disagreement between methods, especially at low carnosine concentrations (figure 6, panel E). This is further confirmed by plotting individual absolute values obtained with 1H-MRS against HPLC, which shows that most data fall far from the identity line (figure 6, panel F).

**Figure 6.**
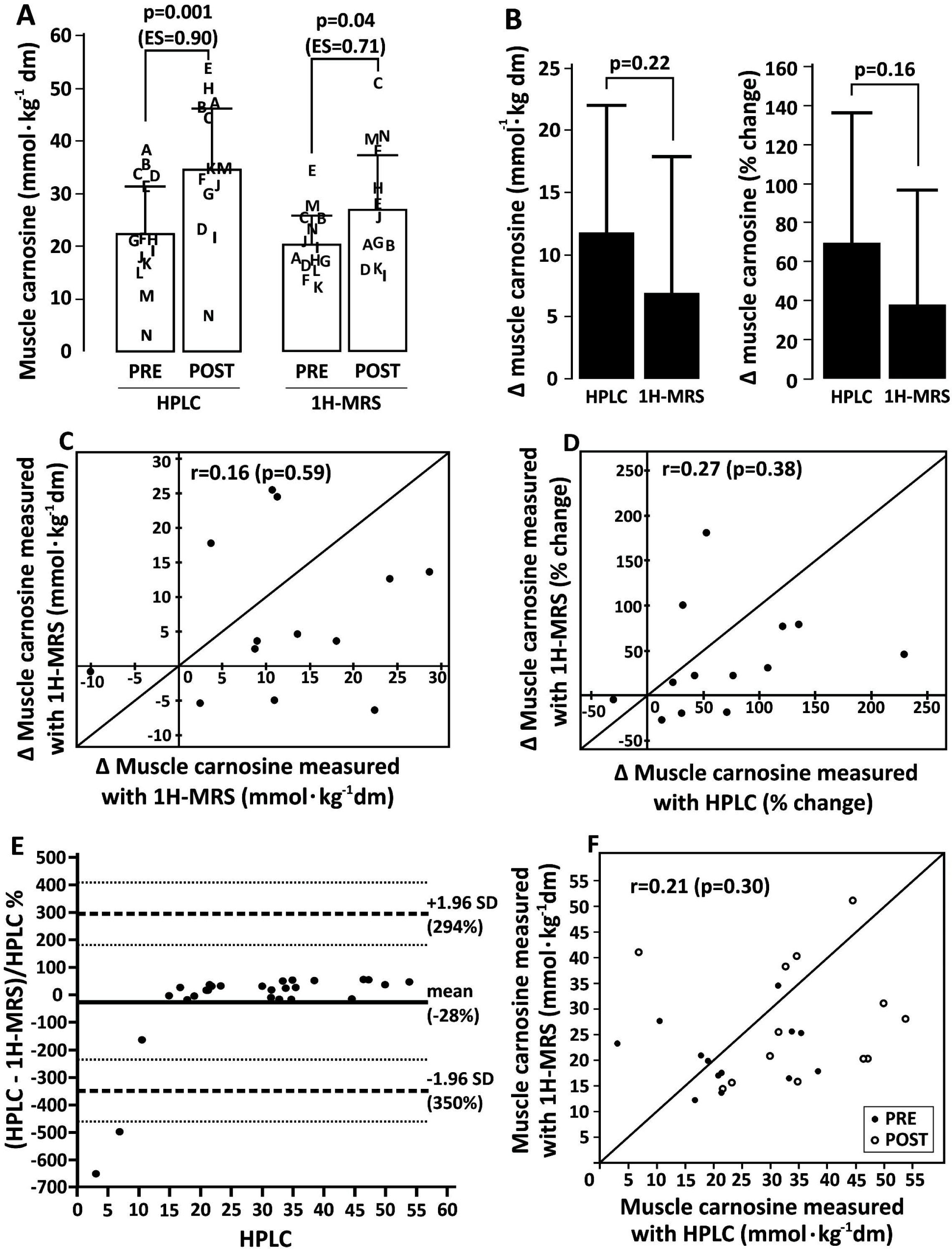
Discriminant validity of 1H-MRS for *in vivo* muscle carnosine quantification, and convergent validity of 1H-MRS *vs.* HPLC for muscle carnosine determination. Panel A: Individual (represented in letters) and mean±SD values for muscle carnosine measured using both methods before and after β-alanine supplementation. Panel B: Absolute (left chart) and relative (right chart) post-pre delta values for muscle carnosine measured using both methods in response to β-alanine supplementation. Panel C: Absolute post-pre delta values obtained using both methods plotted against the identity line (*i.e.*, representing 100% agreement between methods). Panel D: Relative post-pre delta values obtained using both methods plotted against the identity line (*i.e.*, representing 100% agreement between methods). Panel E: Bland-Altman plot for percent differences between methods. Panel F: pooled pre and post data for muscle carnosine values obtained using both methods plotted against the identity line (*i.e.*, representing 100% agreement between methods).

## Discussion

Considering the growing attention that carnosine in muscle has been receiving due to its potential ergogenic and therapeutic properties, quantifying this dipeptide in muscle tissue is becoming an increasingly necessary procedure. As such, the use of an accurate, reliable and sensitive method for carnosine quantification is of the utmost importance. Although non-invasive alternatives for muscle biopsies are unquestionably important, their validity as well as their limitations must be known. Since no study to date has thoroughly assessed 1H-MRS validity for muscle carnosine quantification, we performed a series of *in vitro* and *in vivo* experiments to examine several aspects of 1H-MRS validity (*i.e.*, signal linearity, matrix effect, reliability, discriminant and convergent validity). To be certain that HPLC is a reliable method for muscle carnosine determination and that it could be used as the reference in this study, we also examined its reliability, which showed excellent repeatability in all instances (*i.e.*, intra-assay, inter-extract, and inter-biopsy).

The present investigation revealed important methodological issues that must be considered when interpreting muscle carnosine values obtained *in vivo* by 1H-MRS. Due to the small signal amplitude [14-17], we sought to be certain that the carnosine signal is quantifiable across the entire physiological range, including the expected values for the lowest and highest extremes of human population, such as those reported in vegetarians [9] and bodybuilders [22]. In this regard, 1H-MRS showed excellent linearity when carnosine is quantified in phantoms containing pure carnosine, as well as a clear ability to detect and quantify carnosine even in the lowest range. Excellent *in vitro* linearity has also been shown by Özdemir *et al*. [11]. However, it must be noted that this does not necessarily imply that quantifying the carnosine signal *in vivo* would be equally precise as linear as the for *in vitro* signal. Although the *in vivo* signal has a good signal to noise ratio of (∼13 in our study), it is clearly broader, of lower amplitude, and presented higher baseline noise when compared to the *in vitro* signal (signal to noise ratio ∼18 in our study). Such phenomenon is similar to the matrix effect often seen in analytical methods and can be explained by the potential influence of fat and bone tissues to the carnosine signal [23,24], as well as the large number of compounds with magnetic nuclei present in human muscle. The magnetic interaction among neighbouring, non-equivalent, magnetic nuclei causes a phenomenon known as spin-spin coupling, where the magnetic field generated by each proton interferes with the other one, resulting in splitting and broadening of signal peaks [25]. This phenomenon can be better observed for the C4-H peak, since its position in the imidazole ring and its proximity with the nearest protons makes its signal more susceptible to spin-spin coupling effects [14]. As herein demonstrated, carnosine peaks already present a small amplitude in the 1H-MRS [17]; when these signals are divided or further weakened by the spin-spin coupling, a decrease in signal occurs, resulting in poorer signal-to-noise ratio [21]. In individuals with low muscle carnosine content, increased error is to be expected, since peak amplitude is naturally lower and, therefore, very close to the basal noise. This is supported by the increased disagreement between 1H-MRS and the reference method shown in the Bland-Altman plot when carnosine concentrations are near to the lowest range.

To investigate the potential impact of the imidazole ring present in other molecules (*e.g.*, free imidazole, free histidine, carnosine analogues and histidine residues in proteins) on the carnosine signal detected by 1H-MRS, a series of acquisitions were performed in phantoms containing pure carnosine, imidazole, histidine, and BSA. Although the best signal quality was obtained with carnosine, quantifiable signals were also obtained with imidazole and free histidine. This indicates that small imidazole-containing molecules might constitute a potential source of error, although they are likely of low relevance for the skeletal muscle since they are expressed in very low concentrations in comparison with carnosine [26]. However, this may be problematic when carnosine is to be measured in tissues expressing higher concentrations of small imidazole-containing molecules. Conversely, no signal was obtained with BSA, probably due to its large size (∼66 kDa). Large molecular size leads to slow tumbling and correspondingly short spin-spin relaxation times (T2), resulting in a complex spectrum with broad peaks of low amplitude that do not surpass noise level. Accordingly, 1H-MRS data become unreliable for proteins larger than 30 kDa at room temperature [27]. These results indicate that large imidazole-rich proteins such as haemoglobin do not represent a source of error, although smaller histidine-rich proteins, such as myoglobin (17 kDa, 4.7 mg·g^-1^ wet muscle, 11% histidine) could still contribute to the *in vivo* signal. However, we unfortunately were unable to prepare a phantom with purified myoglobin for further verification. Hence, whether myoglobin constitutes a source of error requires future clarification.

To assess reliability, convergent and discriminant validity of *in vivo* 1H-MRS, a series of analyses were conducted. No significant differences were shown between test and retest values, indicating that 1H-MRS is free of systematic errors and that the large variation observed is due to random error. Importantly, a remarkable increase in test-retest variation (6.6% *vs.* 16.9%) was shown when the retest was performed with the participant being removed from and then relocated to the scanner, which is a more “real-world” representation of studies assessing muscle carnosine before and after an intervention. Such an increase in variation indicates that voxel positioning and shimming are major sources of random error in 1H-MRS. In the present study, all possible measures were taken to ensure that voxel would be positioned in the same location (*i.e.*, the same experienced technician was responsible for voxel positioning in all exams; voxel positioning at “retest” was done with the image depicting voxel position at “test” as a guide). Nonetheless, a large variation was observed between test and retest measurements, indicating that even small differences in voxel position have a large impact on the results. Such a large variation arising from voxel repositioning and re-shimming is somewhat expected for a metabolite with a broad signal and of low amplitude, such as carnosine, since similar levels of variation have been reported for other metabolites of much sharper and high amplitude signals [28]. In addition, the ∼17% variation reported in this study is not too dissimilar to previous investigations [11, 13, 17].

One explanation for the larger variation of 1H-MRS is the non-homogeneity of carnosine distribution in the skeletal muscle. Carnosine content is greater in type II than type I muscle fibres [9]; therefore, the inevitable change in muscle site when performing two 1H-MRS, albeit small, may cause sampling sites to have different fiber type composition, adding a source of measurement error. However, slight changes in sampling sites likely occurred with muscle biopsies and, yet, the variation for HPLC was lower. This means that other factors may play a role in the increased variability in 1H-MRS, such as the proximity with tissues that may cause signal interference (*e.g.*, adipose tissue). The fat signal often appears bright in many important clinical imaging sequences and can obscure other signals [29, 30]. Thus, adipose tissue near or at the data acquisition site can contribute to the increased variability. Additionally, 1H-MRS appears to be more sensitive to changes in sampling sites because shimming and spin-spin coupling are dependent on the angle between spins and the magnetic field, which may alter with slight changes in sampling sites. Such orientation-dependence is particularly true for the carnosine signal in human skeletal muscle [31]. Participants’ motion during the exam could also disrupt data acquisition, thereby contributing to 1H-MRS variability [32], although all participants were requested to remain still during the entire duration of the exams.

Another putative contributing factor for the poor reliability of 1H-MRS in our study is the spectral correction that is applied as default in the Phillips post-acquisition data processing algorithm (Dr Wim Derave, personal communication). Developed to optimise the spectrum obtained brain, the default mode (on) herein used corrects the metabolite spectra using the water-unsuppressed spectra. However, if fat tissue is present in the voxel region, the baseline can be distorted, possibly affecting the carnosine signal. Although this has never been reported in the literature, we tested whether changing the spectral mode to “off” impacts carnosine signal. Interestingly, no significant differences were observed when the spectra obtained with the correction “on” were reconstructed to the mode “off” (n=11 spectra obtained from 5 young men; p=0.69 mode “on” *vs.* mode “off”). Moreover, we observed no improvement in test-retest reliability (removing and repositioning the participants in the scanner) by changing the spectral correction mode to “off” (CV=36.8%, n=3). Thus, it is unlikely that the results herein presented are artifactual findings specific to our acquisition settings, and they probably reflect true limitations of the 1H-MRS to quantify carnosine in skeletal muscle.

The discriminant validity study showed that 1H-MRS, despite having large variation owing to random errors, is sensitive to detect group-mean increases in muscle carnosine in response to β-alanine supplementation. The changes shown with 1H-MRS are similar to those reported in other studies [33]. Yet, when comparing the ability of 1H-MRS to detect changes in muscle carnosine with HPLC, a high degree of disagreement between methods was shown. Likewise, a large degree of disagreement was shown in all instances where 1H-MRS and HPLC were compared, which becomes particularly evident when the results are analysed at the individual level (Figure 6). The disagreement seemed to increase when participants showed smaller carnosine content, possibly due to 1H-MRS signal characteristics. As discussed, the carnosine peak amplitude in 1H-MRS makes it prone to suffer noise interference, which is more pronounced at lower concentrations. On the other hand, carnosine peaks in the HPLC chromatogram are large, sharp and easily quantifiable across the entire physiological range. Previous studies in the literature also appear to support the discrepancy between 1H-MRS and HPLC measurements of muscle carnosine, since HPLC studies consistently show 40-80% increases in muscle carnosine in response to 4 weeks of a 6.4 g.day^-1^ dosage of β-alanine supplementation [8,9,19], whereas studies using 1H-MRS show much larger increases of 140-160% in muscle carnosine [12], or no increase at all [13] following similar β-alanine supplementation protocols. In line with this, a recent meta-analysis including numerous studies assessing the effects of β-alanine supplementation on muscle carnosine showed that the method (HPLC *vs.* 1H-MRS) is a factor that significantly affects muscle carnosine outcomes (Resende *et al*., submitted). It seems that the strength of the magnetic field of the scanner could be an important factor affecting the validity of muscle carnosine measurements. While the sole study unable to detect increases in muscle carnosine following β-alanine supplementation used a 1.5T scanner [13], improved reliability was shown with carnosine being quantified with a 7T scanner [17].

A potential limitation of this study is that the results were obtained from the *m. gastrocnemius medialis*, known to have a mixed proportion of type I and II fibers. Hence, it remains to be elucidated whether 1H-MRS shows better measurement performance for carnosine quantification in a more homogeneous muscle. In addition, considering that physically active participants were recruited for the present study and that local adipose tissue appeared to be an important factor interfering with 1H-MRS signal [23], we cannot rule out the possibility that 1H-MRS is more reliable in individuals with lower body fat, such as athletes.

To conclude, 1H-MRS is capable of measuring carnosine in muscle tissue and is sensitive to detect overall changes in muscle carnosine brought about by β-alanine supplementation. However, 1H-MRS has a high degree of variation due to random error associated with a large effect that minor variations in voxel positioning has on the carnosine signal. This also results in poor convergent validity. This makes quantification problematic, particularly in regions surrounded by fat tissue, in individuals with high levels of body fat, and in individuals with low muscle carnosine levels. Caution should be exercised when interpreting muscle carnosine quantification data obtained with 3-T 1H-MRS, especially when absolute quantification is used. Future studies assessing muscle carnosine non-invasively should opt for scanners with stronger magnetic fields, such as 7-T 1H-MRS, for improved resolution.

## Methods

### Ethical approval

The study was approved by the Ethics Committee of the School of Medicine of the University of Sao Paulo (approval number – 43986415.3.0000.0065) and conformed to the 2013 version of the Declaration of Helsinki. All participants signed the written informed consent term before taking part of the study.

### *In vitro* investigation

Thirteen 0.5 L cylindrical bottles mimicking a human calf were filled with solidified solutions of carnosine, imidazole, histidine or BSA. Phantoms of 6 different carnosine concentrations (3.0, 4.5, 6.0, 12.5, 25.0 and 50.0 mmol·L^-1^) were prepared to assess signal linearity within the physiological range and signal behaviour near to the lowest physiological range. Phantoms of 3 different imidazole and histidine concentrations (12.5, 25.0 and 50.0 mmol·L^-1^ each) were prepared to examine whether the signals would differ between imidazole-containing substances. One phantom containing BSA (the equivalent to 12.5 mmol·L^-1^ of imidazole) was prepared to assess whether imidazole-residues in large size molecules (*e.g.*, protein) could emit a signal at the same chemical shift (7 and 8 ppm). All phantoms were solidified by melting agarose 2% w:v in autoclaved ultra-pure water prior to adding carnosine, imidazole, histidine or BSA. Phantom concentrations were calculate based on the imidazole content so that all concentrations were equimolar to 12.5 mmol·L^-1^ of imidazole. The 12.5 mmol·L^-1^ concentration for BSA was chosen because this is nearly the maximum achievable within the solubility of BSA and it represents a mid-range physiological concentration of carnosine in human skeletal muscle.

For the present investigation, a 3 Tesla whole-body magnetic resonance scanner (Achieva, Philips, Best, The Netherlands) equipped with an 8-channel knee coil was used. All the spectra were acquired using single voxel point-resolved spectroscopy (PRESS) localization with the following parameters: TR/TE=6000/30 ms, voxel size=10×10×10 mm^3^, number of averages (NEX)=224, 2048 data points with a spectral width of 2000 Hz. The total acquisition time was 20 min for each phantom. All spectra were processed in jMRUI software. Residual water and lipid peaks were removed by a Hankel Lanczos Squares Singular Values Decomposition (HSVLD) algorithm from the carnosine, histidine and BSA spectra, and their C2-H and C4-H peaks were fitted with Advanced Method for Accurate, Robust and Efficient Spectral fitting (AMARES) using single Lorentzian line shapes. Carnosine’s signal linearity was evaluated by linear regression of the “carnosine concentration *vs*. signal” calibration curve.

### *In vivo* investigation

Sixteen young, healthy, physically active men volunteered to participate, two of whom could not complete the entire study due to personal reasons. Therefore, 14 participants (age:27±5 years; body mass: 82.9±11.8 kg; stature: 1.77±0.06 m; body mass index: 26.3±2.4 kg·m^2^) completed all tests. Participants were fully informed of possible risks and discomforts associated with participation. They were requested to maintain similar levels of physical activity and dietary patterns for the duration of the study. Exclusion criteria were: *i)* use of supplements containing creatine or β-alanine in the 3 months and 6 months prior to the study; *ii)* use of anabolic steroids; *iii)* chronic use of glucocorticoids; *iv)* chronic-degenerative disease and/or condition that affected the locomotor apparatus, and *v)* any condition that would prevent them from undertaking the proposed tests (*e.g.*, metallic prostheses that could interfere with 1H-MRS quality).

Participants were assessed for muscle carnosine before and after a 4-week period of β-alanine supplementation. To assess convergent validity, muscle samples were obtained from the same group of participants immediately after 1H-MRS, both before and after β-alanine supplementation, so that the results obtained with 1H-MRS could be compared with those obtained with HPLC. Muscle carnosine concentrations obtained with 1H-MRS were converted to the same unit of muscle carnosine content (*i.e.*, from mmol·L^-1^ to mmol·kg^-1^ of dry tissue) for a clearer comparison between methods. The 1H-MRS *in vivo* and muscle biopsy assessments were individually standardized so that each participant performed their PRE and POST-sessions at the same time of day. Participants were requested to abstain from alcohol and unaccustomed exercise in the 48 hours prior to the experimental sessions. Participants were instructed to arrive at the laboratory at least 2 hours following their last meal. *Ad libitum* water consumption was allowed before and after the sessions.

β-alanine was provided in 800-mg tablets (CarnoSyn™, NAI, USA) and the participants were asked to take 2 tablets along with meals, four times per day, totalling 6.4g·d^-1^ of β-alanine. This intervention was intentionally chosen due to its highly consistent effects on muscle carnosine [2, 8]. All 16 participants completed a baseline assessment for carnosine quantification in the medial portion of *m. gastrocnemius* of the right leg using both 1H-MRS and HPLC. Carnosine quantification via 1H-MRS was not possible in one participant due to a peak of very small amplitude with baseline below zero. Therefore, the analysis of convergent validity was conducted on 15 participants. To minimize differences between methods owing to variations in sampling sites, biopsy sites were intentionally taken from the closest possible sites to those where the spectra were obtained. This was ensured with the physician examining the image of the voxel position before defining the location and the depth the biopsy needle would be inserted. Following the supplementation period, the 14 participants who completed the entire study were again assessed for muscle carnosine using both 1H-MRS and HPLC. The responses to supplementation were used to compare the discriminant validity between methods.

A sub-sample of 10 participants volunteered for the test-retest reliability assessment of 1H-MRS with voxel repositioning and re-shimming. They undertook the first 1H-MRS, left the room, waited for 5 minutes and then were repositioned back on the machine for the second 1H-MRS. The voxel was repositioned as closely as possible to the site where it was positioned in first test; this was achieved using an image of the voxel position obtained in the first 1H-MRS as a guide. Another sub-sample of 5 participants volunteered for the test-retest reliability assessment of 1H-MRS without voxel repositioning and re-shimming. They undertook the first 1H-MRS and stood still for the second 1H-MRS, which was performed immediately after the first.

To assess intra-assay reliability of HPLC, muscle extracts obtained from 15 biopsy samples were analysed in duplicate in two independent runs. To assess “inter-extract” reliability of HPLC, two different muscle extracts obtained from 11 biopsy samples (*m. vastus lateralis*) randomly chosen from this same collection were analysed in duplicates on different days. To assess “inter-biopsy” reliability of HPLC, two consecutive muscle samples (*m. gastrocnemius*) were obtained from the same incision in a sub-sample of 7 participants who volunteered for this study. The second biopsy location was changed by rotating the needle’s window guillotine in 90 degrees, thereby sampling the collateral site of the first biopsy.

### 1H-MRS *in vivo* assessment

Spectra were acquired using single voxel point-resolved spectroscopy (PRESS) localization with the following parameters: TR/TE=3000/30 ms, voxel size=10×10×30 mm^3^, number of averages (NEX)=256, 2048 data points with a spectral width of 2000 Hz. The total acquisition time of the 1H-MRS was 13.9 min. The 50 mmol·L^-1^ carnosine phantom was used as an external reference. To that end, an acquisition was made using the same parameters as those used *in vivo*, except for the TR, which was 12000 ms. In each *in vivo* measurement, the right leg of each participant was positioned and was firmly immobilized in the knee coil, such that the gastrocnemius muscle was in the centre of the coil. The left leg was supported outside the coil to improve comfort and thus minimize leg movement. Voxel location was standardized on the larger calf region in the centre of the medial portion of the gastrocnemius muscle of the right leg. The same well-trained and experienced biomedical technician was responsible for placing the voxel in all conditions. After placing the voxel, a set of images depicting individual voxel location was saved and used to guide positioning in all further exams of that individual.

### Quantification

Absolute quantification of the carnosine resonance was determined using the following equation [21]:

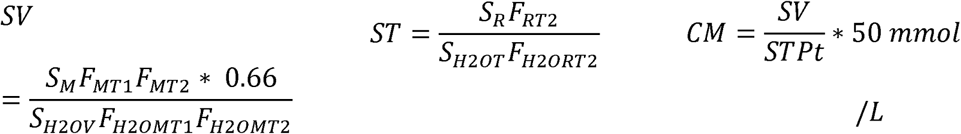

CM is the concentration of the metabolite *in vivo*, SV and ST are the signals of the water-corrected metabolite *in vivo* and *in vitro*; SM is the integral of the carnosine peak *in vivo* and SR is the integral of the carnosine peak *in vitro*; F_MT1_ is the correction factor for T1 relaxation of the metabolite *in vivo*; F_MT2_ and F_RT2_ are the correction factors for metabolic T2 relaxation *in vivo* and *in vitro*; S_H2OV_ is the integral of the water peak *in vivo* and S_H2OT_ is the integral of the water peak *in vitro*; F_H2OMT1_ is the correction factor for T1 relaxation of water *in vivo*; And F_H2OMT2_ and F_H2ORT2_ are the correction factors for T2 relaxation of water *in vivo* and *in vitro*. Pt is the temperature correction factor applied as the signal decreases by 6% between the room temperature (*i.e.*, phantom temperature) and body temperature [34]. For the *in vitro* signal, it is not necessary to correct the T1 relaxation, since the acquisition was performed with a sufficiently long TR (TR=12000 ms) to neglect this factor. Signals were also corrected by water content; since the water content in phantoms is ∼100%, a correction factor=1 was used. For the *in vivo* analyses, a correction factor=0.66 was used, assuming that ∼2/3 of the muscle is water [35].

The relaxation correction factors were calculated using the following equations:

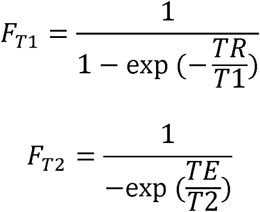

T1 and T2 values for water and carnosine were taken from the literature, and were assumed to be 1420 ms and 32 ms for water in muscle [36] and 520 ms and 66 ms for *in vivo carnosine* [11]. The T2 values of *in vitro* water and carnosine *in vitro* (52 ms and 200 ms) were measured using different TEs (31, 61, 99, 150, 228, and 400 ms) with TR of 6000 ms and calculated using MATLAB^®^ software (MathWorks, Natick, MA, USA) by fitting data points determined using the peak areas of metabolites to a mono-exponential function Ms=M0 exp(-TE/T2). To further convert concentration values in mmol·L^-1^ into the content equivalent in mmol·kg^-1^ of dry muscle, results were multiplied by 3.3, a factor which assumes that for every 1 kg of dry muscle there is 3.3 kg of water. Spectrum figures are presented with the raw (untreated) spectrum on the left side and a cut window (framework of the target the frequency) on the right. The cut window signal was normalized by the baseline offset to facilitate visual comparison between both situations [36].

### Muscle biopsies

Muscle samples (∼70-100 mg) were obtained under local anaesthesia (3 mL, 1% lidocaine) from the mid-portion of the *m. gastrocnemius* using the percutaneous needle biopsy technique with suction [37]. Samples were obtained from the same leg for all experiments. PRE and POST supplementation biopsies were taken from incisions made as close as possible to one another. Samples taken for the inter-biopsy reliability analyses were obtained from the same incision, but from slightly different sites, as described above. All samples were snap frozen in liquid nitrogen and were subsequently stored at −80°C until analyses. Samples were freeze-dried and dissected free of any visible blood, fat and connective tissue before being powdered and further submitted to HPLC determination of carnosine.

### Chromatographic determination of histidine-containing dipeptides in whole muscle

Deproteinized muscle extracts were obtained from 3-5 mg freeze-dried samples according to the protocol described by Harris *et al*. [8]. Briefly, samples were deproteinized with 0.5M HClO_4_ and neutralized with 2.1M KHCO_3_. The extracts were then filtered through a 0.22 μm centrifugal PVDF filter unit and stored at −80^°^ C until analysis. Total muscle carnosine content was quantified by HPLC (Hitachi, Hitachi Ltd., Tokyo, Japan), according to the method described by Mora *et al* [38]. Mobile phases consisted of solvent A, containing 0.65 mM ammonium acetate, pH 5.5, in water/acetonitrile (25:75); and solvent B, containing 4.55 mM ammonium acetate, pH 5.5, in water/ acetonitrile (70:30). The chromatographic separation was developed using an Atlantis HILIC silica column (4.6 × 150 mm, 3 μm; Waters, Milford, MA, USA) attached to an Atlantis Silica column guard (4.6 x 20 mm, 3 μm), at room temperature. The analysis conditions comprised of a linear gradient from 0 to 100% of solvent B in 13 min at a flow rate of 1.4 mL·min^-1^. Separation was monitored using a U.V. coupled detector at a wavelength of 214 nm.

### Statistical analyses

Signal linearity of the *in vitro* analysis was verified by interpolating signal intensity by concentration and calculating Pearson correlation coefficient (r). Test-retest reliability was assessed using: *1)* a paired sample *t*-test to check for systematic errors, *2)* intraclass correlation coefficient (ICC - two-way random, absolute agreement, single-measures) with their respective 95% confidence intervals (95%CI) and *3)* within-subject mean square root coefficient of variation (CV) [39]. Test-retest data were plotted to individually display the distance between data pairs and the identity line. Convergent validity was assessed using: *1)* an unpaired sample *t*-test to check for differences between HPLC and 1H-MRS values, for both PRE and POST supplementation data sets, and *2)* the Bland-Altman plot for percentage differences against mean values for the overall data set. HPLC *vs*. 1H-MRS data was plotted to individually display the distance between data pair and the identity line. Discriminant validity was assessed using a paired sample *t*-test to compare mean carnosine values between PRE and POST supplementation period. Effect sizes (ES) for the muscle carnosine content increase after β-alanine supplementation were calculated using Cohen’s *d*. All analyses were conducted in the IBM SPSS software (version 20). Bland-Altman was built using the GraphPad Prism software (version 5.03).

## Acknowledgements

The authors would like to thank Dr Leslie Boobis for the very generous medical training on muscle biopsies. We also thank Felipe Gregório Jardim, Mariana Franchi and Kleiner Márcio de Andrade Nemezio for the help during data collection and Rodrigo Silva Maeda for the help during analysis stage. Vinicius da Eira Silva was financially supported by Coordenação de Aperfeiçoamento de Pessoal de Nível Superior. Bruno Gualano, Vitor de Salles Painelli and Guilherme Giannini Artiol have been financially supported by Fundação de Amparo à Pesquisa do Estado de Sao Paulo (FAPESP grant numbers: 2013/14746-4, 2013/04806-0 and 2014/11948-8). This study was supported by CAPES-PROEX. This study was financed in part by the Coordenação de Aperfeiçoamento de Pessoal de Nível Superior - Brasil (CAPES) - Finance Code 001.

## Author Contributions Statement

Conceptualization V.E.S, G.G.A, C.S, B.G, V.S.P and M.C.O; Methodology, V.E.S, G.G.A, and M.C.O.; Investigation, V.E.S, G.G.A, V.S.P., M.C.O, S.K.S, W.R.P; Formal Analysis V.E.S, G.G.A, and M.C.O..; Writing – Original Draft, V.E.S, G.G.A, V.S.P; Writing – Reviewing & Editing, V.E.S, G.G.A, V.S.P., M.C.O, S.K.S, C.S, B.G, E.M.C, W.R.P.

## References

1. Boldyrev, A.A., Aldini, G. & Derave, W. Physiology and pathophysiology of carnosine. Physiol Rev. 93, 1803–1845 (2013).

2. Saunders, B. et al. β-alanine supplementation to improve exercise capacity and performance: a systematic review and meta-analysis. Br J Sports Med. 51, 658–669 (2017).

3. Artioli, G.G., Sale, C. & Jones, R.L. Carnosine in health and disease. Eur J Sport Sci. 19, 30–39 (2018).

4. Matthews, J.J., Artioli, G.G., Turner, M.D., Sale C. The Physiological Roles of Carnosine and β-Alanine in Exercising Human Skeletal Muscle. Med Sci Sports Exerc. 51, 2098–2108 (2019).

5. Dolan, E. et al. A Comparative Study of Hummingbirds and Chickens Provides Mechanistic Insight on the Histidine Containing Dipeptide Role in Skeletal Muscle Metabolism. Sci Rep. 8, 14788, 10.1038/s41598-018-32636-3 (2018).

6. Carvalho, V.H. et al. Exercise and β-alanine supplementation on carnosine-acrolein adduct in skeletal muscle. Redox Biol, 18, 222–228 (2018).

7. Ghodsi, R. & Kheirouri, S. Carnosine and advanced glycation end products: a systematic review. Amino Acids. 50, 1177–1186 (2018).

8. Harris, R.C. et al. The absorption of orally supplied beta-alanine and its effect on muscle carnosine synthesis in human vastus lateralis. Amino Acids. 30, 279–289 (2006).

9. De Salles Painelli, V. et al. High-Intensity Interval Training Augments Muscle Carnosine in the Absence of Dietary Beta-alanine Intake. Med Sci Sports Exerc. 50, 2242–2252 (2018).

10. Neves, M. Jr et al. Incidence of adverse events associated with percutaneous muscular biopsy among healthy and diseased subjects. Scand J Med Sci Sports. 22, 175–178 (2012).

11. Ozdemir, M.S. et al. Absolute quantification of carnosine in human calf muscle by proton magnetic resonance spectroscopy. Phys Med Biol. 52, 6781–6794 (2007).

12. Chung, W., Baguet, A., Bex, T., Bishop, D.J. & Derave, W. Doubling of muscle carnosine concentration does not improve laboratory 1-hr cycling time-trial performance. Int J Sport Nutr Exerc Metab. 24, 315–324 (2014).

13. Black, M.I. et al. The Effects of β-Alanine Supplementation on Muscle pH and the Power-Duration Relationship during High-Intensity Exercise. Front Physiol. 9, 111, 10.3389/fphys.2018.00111 (2018).

14. Boesch, C. & Kreis, R. Dipolar coupling and ordering effects observed in magnetic resonance spectra of skeletal muscle. NMR Biomedicine. 14, 140–148 (2001).

15. Kreis, R. Quantitative localized 1H MR spectroscopy for clinical use. Prog NMR Spectrosc. 31, 155–195 (1997).

16. Tkac, I., Ugurbil, K. & Gruetter, R. On the quantification of low concentration metabolites by 1H NMR spectroscopy in the human brain at 7 Tesla. Proceedings of the 10th Meeting of the International Society of Magnetic Resonance in Medicine, Honolulu, USA. 528 (2002).

17. Just Kukurová, I. et al. Improved spectral resolution and high reliability of in vivo 1H MRS at 7 T allow the characterization of the effect of acute exercise on carnosine in skeletal muscle. NMR Biomed. 29, 24–32 (2016).

18. Solis, M.Y. et al. Effects of beta-alanine supplementation on brain homocarnosine/carnosine signal and cognitive function: an exploratory study. PLoS One, 10, e0123857. 10.1371/journal.pone.0123857 (2015).

19. Hill, C.A. et al. Influence of beta-alanine supplementation on skeletal muscle carnosine concentrations and high intensity cycling capacity. Amino Acids. 32, 225–233 (2007).

20. Alkemade, C.T.J. et al. A review and tutorial discussion of noise and signal-to-noise ratios in analytical spectrometry — I. Fundamental principles of signal-to-noise ratios. Spectrochim Acta Part B At Spectrosc. 33, 383–399 (1978).

21. Hoult, D.I. & Richards, R.E. The signal-to-noise ratio of the nuclear magnetic resonance experiment. J Magn Reson. 1, 71–85 (1969).

22. Tallon, M.J., Harris, R.C., Boobis, L.H., Fallowfield, J.L. & Wise, J.A. The carnosine content of vastus lateralis is elevated in resistance-trained bodybuilders. J Strength Cond Res. 19 (2005).

23. Mon, A., Abé, C., Durazzo, T.C. & Meyerhoff, D.J. Potential effects of fat on magnetic resonance signal intensity and derived brain tissue volumes. Obes Res Clin Pract. 10, 211–215 (2016).

24. Mon, A., Abé, C., Durazzo, T.C. & Meyerhoff, D.J. Effects of fat on MR measured metabolite signal strengths: implications for in vivo MRS studies of the human brain. NMR Biomed. 26, 1768–1774 (2013)

25. Due, C.O., Weber, O.M., Trabesinger, A.H., Meier, D. & Boesiger, P. Quantitative 1H MRS of the human brain in vivo based on the simulation phantom calibration strategy. Magn Reson Med. 39, 491–496 (1998).

26. Parkhouse, W.S., McKenzie, D.C., Hochachka, P.W. & Ovalle, W.K. Buffering capacity of deproteinized human vastus lateralis muscle. J Appl Physiol. 58, 14–17 (1985).

27. Wand, A.J., Ehrhardt, M.R. & Flynn, P.F. High-resolution NMR of encapsulated proteins dissolved in low-viscosity fluids. Proc Natl Acad Sci. 95, 15299–15302 (1998).

28. Al-iedani, O., Lechner-Scott, J., Ribbons, K. & Ramadan, S. Fast magnetic resonance spectroscopic imaging techniques in human brain-applications in multiple sclerosis. J Biomed Science. 24, 17 (2017).

29. Bley, T.A., Wieben, O., François, C.J., Brittain, J.H. & Reeder, S.B. Fat and water magnetic resonance imaging. J Magn Reson Imag. 31, 4–18 (2010).

30. Maudsley, A.A., Govind, V. & Arheart, K.L. Associations of age, gender and body mass with 1H MR-observed brain metabolites and tissue distributions. NMR Biomed. 25, 580–593 (2012).

31. Kreis, R. & Boesch, C. Orientation dependence is the rule, not the exception in 1H-MR spectra of skeletal muscle: the case of carnosine. ISMRM Proc Intl Soc Magn Reson Med. 31 (2000).

32. Marshall, I., Wardlaw, J., Cannon, J., Slattery, J. & Sellar, R.J. Reproducibility of metabolite peak areas in 1H MRS of brain. Magn Reson Imaging. 14, 281–292 (1996).

33. Derave, W. et al. Beta-alanine supplementation augments muscle carnosine content and attenuates fatigue during repeated isokinetic contraction bouts in trained sprinters. J Appl Physiol. 103, 1736–1743 (2007).

34. Davies, G.R. et al. Preliminary magnetic resonance study of the macromolecular proton fraction in white matter: a potential marker of myelin? Mult Scler. 9, 246–249 (2003).

35. Schoeller, D.A. Changes in total body water with age. Am J Clin Nutr. 50, 1176–1181, DOI: 10.1093/ajcn/50.5.1176 (1989).

36. Kohl, S.M. et al. State-of-the art data normalization methods improve NMR-based metabolomic analysis. Metabolomics. 8, 146–160 (2012).

37. Bergstrom, J. Muscle electrolytes in man determination by neutron activation analysis on needle biopsy specimens. A study on normal subjects, kidney patients and patients with chronic diarrhea. Scand J Clin Lab Invest. 14, 100–110 (1962).

38. Mora, L., Sentandreu, M.A. & Toldrá, F. Hydrophilic chromatographic determination of carnosine, anserine, balenine, creatine, and creatinine. J Agric Food Chem. 55, 4664–4669 (2007).

39. Hyslop, N.P. & White, W.H. Estimating precision using duplicate measurements. J Air Waste Manag Assoc. 59, 1032–1039 (2009).

